# SUN2 mediates epigenetic remodeling to drive mechanotransduction during skin fibrosis

**DOI:** 10.64898/2026.03.19.712957

**Authors:** Aya Nassereddine, Kerri Davidson, Sandra Sandria, Jyot D. Antani, Shreyasi Das, Rachel Rivera, Van Anh Tran, Monique Hinchcliff, Megan King, Valerie Horsley

## Abstract

Fibrosis involves sustained changes in fibroblast gene expression, leading to excessive extracellular matrix (ECM) deposition and progressive tissue stiffening. Although matrix stiffness is a potent regulator of cell fate and transcription, it is not clear how nuclear mechanosensing contributes to fibrosis. Here, we define a central role for SUN2, a component of linker of nucleoskeleton and cytoskeleton (LINC) complexes, as a mediator of stiffness-dependent nuclear and chromatin responses during skin fibrosis. *SUN2* transcripts are upregulated in dermal fibroblasts of patients with systemic sclerosis and Sun2 protein is elevated in fibrotic mouse skin. Nuclear size, A-type lamins and Sun2 are elevated in dermal fibroblasts plated on stiff substrates. Loss of *Sun2* protects against bleomycin-induced skin fibrosis in vivo and abolishes stiffness-induced changes in nuclear size and fibrotic gene expression in vitro. Mechanistically, we identify three *Sun2*-dependent mechanosensitive chromatin states and show that mechanical induction of the histone methyltransferase Ezh2 requires *Sun2*. These findings define SUN2 as a nuclear mechanosensor that couples matrix stiffness to chromatin regulation and transcriptional programs that drive fibrosis, identifying it as a potential therapeutic target pathway in fibrotic disease.

## Introduction

Fibrosis, defined as a pathologic accumulation of extracellular matrix (ECM) proteins, is believed to cause 45% of all deaths in the industrialized world, but it has few effective therapies ***Wynn (2004***). Fibroblasts produce and remodel the ECM, and in fibrosis this process leads to stiffening that disrupts normal tissue function. Fibroblast-induced tissue stiffening provides a feed-forward loop to amplify fibrosis, in part by promoting a persistent “memory” state in which fibroblasts remain activated even in the absence of pro-fibrotic mechanical cues from the matrix driven in part by epigenetic changes ***Balestrini et al. (2012***); ***Caliari et al. (2016); Liu et al. (2010***). While mechanical stimuli, such as ECM stiffening during fibrosis, are key regulators of cell migration, proliferation, and differentiation ***Engler et al. (2006); Jaalouk and Lammerding (2009); Jansen et al. (2017); Fritzsche (2021***), how mechanosensing regulates fibrosis is not fully understood.

One potential mechanism that links mechanosensing to fibrosis is nuclear mechanotransduction. The shape, polarization, and positioning of the nucleus are controlled by external mechanical forces that contract cytoskeletal filaments and can be directly translated to the nuclear membrane and nucleoplasmic lamins via linker of nucleoskeleton and cytoskeleton (LINC) complexes ***Smith et al. (2022); Chang et al. (2015a); Lombardi and Lammerding (2011); Luxton et al. (2011); Starr (2010); Carley et al. (2021***). Stiff substrates can induce tension on LINC complexes and alter chromatin accessibility and fibrotic memory ***Walker et al. (2021***). The LINC complex is composed of SUN (Sad1/UNC-84)-domain containing proteins that span the inner nuclear membrane and KASH (Klarsicht/ANC-1/Syne Homology)-domain containing proteins in the outer nuclear membrane that interact in the cytoskeleton ***King (2023***). Interestingly, fibroblasts that express a dominant negative KASH protein can shift to a non-contractile morpholgy when moved from stiff to soft substrates, indicating a role in myofibroblast mechanosensing “memory” ***Walker et al. (2021***), and our previous work showed that mice lacking *Sun2* display reduced cardiac fibrosis after induction of cardiac hypertrophy ***Stewart et al. (2019***). Furthermore, overexpression of a dominant negative SUN domain construct in fibroblasts did not have major effects on ECM gene expression or nuclear morphology changes associated with substrate stiffness ***Alam et al. (2016***). Thus, mechanisms by which nuclear mechanosensing functions in fibrosis, especially in vivo, are not fully understood.

The mammalian skin is an excellent model to define the roles of nuclear mechanosensing in fibrosis. Mutations in A-type lamin proteins that interact with LINC complexes cause lipodystrophy and induction of collagen production, which are phenotypes associated with skin fibrosis in human patients and mouse models ***Nguyen et al. (2007); Béréziat et al. (2011); Hinz (2012); Tschumperlin et al. (2018); Caves et al. (2025***). For instance, dermal adipocytes undergo lipodystrophy and dermal fibroblasts increase ECM production and remodeling during skin fibrosis***Caves et al. (2025); Varga and Marangoni (2017***). However, whether LINC-mediated nuclear mechanosensing actively drives chromatin remodeling and transcription in fibrosis is not known.

Here, we investigate the impact of nuclear mechanosensing on skin fibrosis. We find that skin fibrosis is associated with altered nuclear architecture and elevated expression of A-type lamins and the LINC complex protein SUN2 in human patients and mouse models of bleomycin induced fibrosis. We find that SUN2-dependent nuclear mechanotransduction tunes fibrotic gene expression and chromatin accessibility to regulate fibrotic responses in both mouse models in vivo and in response to mechanical stimuli in vitro. These data highlight the importance of nuclear mechanosensing in skin fibrosis and reveal that LINC complexes can direct changes in chromatin and gene expression essential for ECM homeostasis.

## Results

### *SUN2* expression is elevated in skin fibroblasts of patients with systemic sclerosis and mice treated with bleomycin

We were intrigued that the inner nuclear membrane protein *SUN2* was differentially regulated in SFRP2+ progenitor fibroblasts in the skin of people without SSc compared to the skin of patients with SSc ***Tabib et al. (2021***). To further examine whether *Sun2* mRNA is altered in fibroblasts in cutaneous fibrosis, we analyzed *SUN2* transcript levels in single-cell RNA sequencing data (scRNA seq) from skin samples of patients without SSc (Normal) and people with SSc (***Odell et al. (2022***), GEO: GSE214088)(Fig. 1A). Analysis of *Sun2* expression in Normal vs SSc fibroblasts revealed a highly significant upregulation of *Sun2* in cells from patients with SSc (Fig. 1B). We confirmed that this shift was consistent across the disease group and not driven by outliers, suggesting SUN2 is a broadly relevant marker of the SSc fibroblast phenotype.

**Figure 1.**
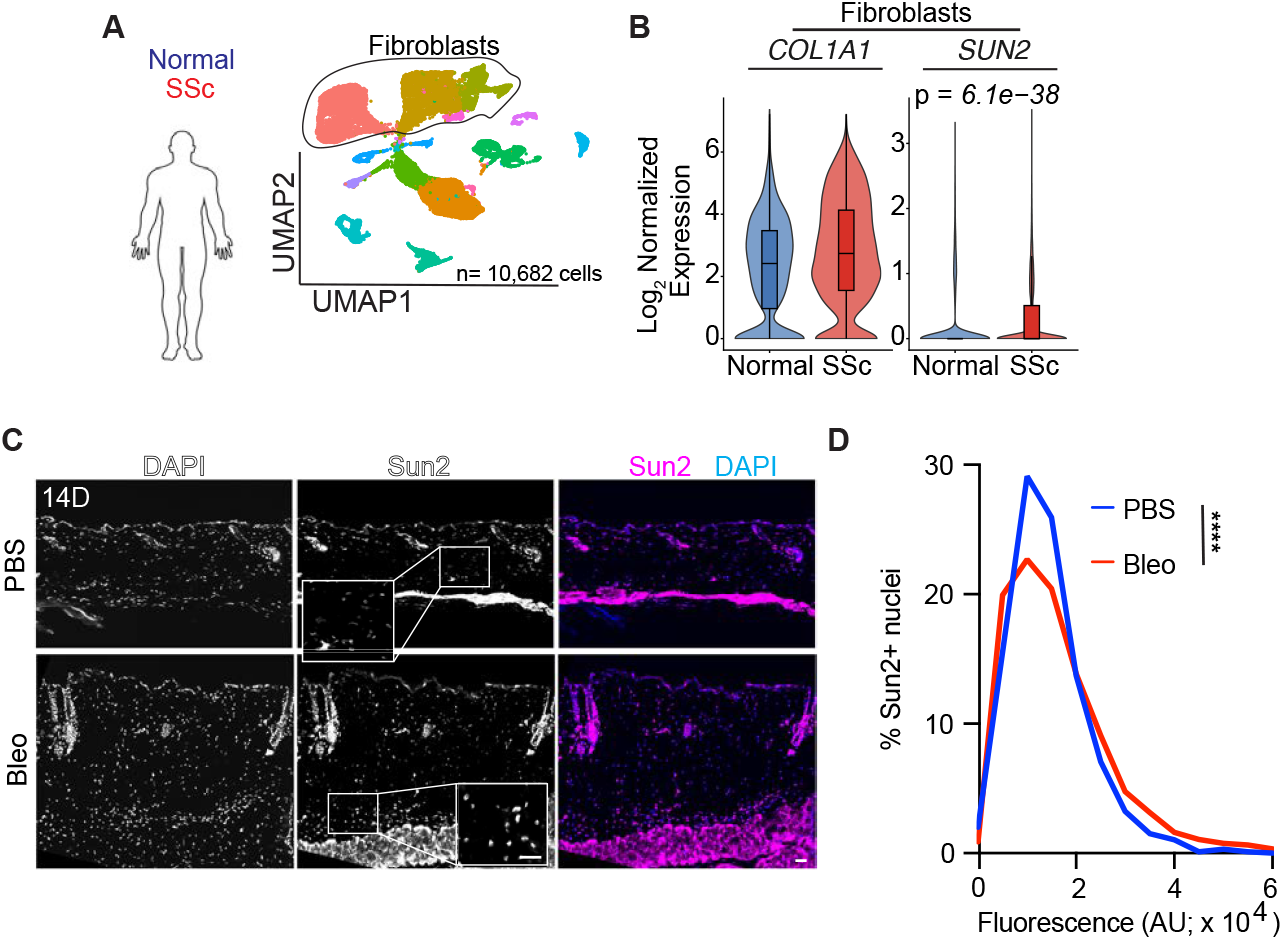
SUN2 increases in fibrotic skin. (A) UMAPs of cell types in skin from 5 patients without (Normal) and 5 patients with Systemic Sclerosis (SSc). GEO: GSE214088. B) Violin plots showing log-normalized expression of *SUN2* and *COL1A1* in dermal fibroblasts. n= 10,682 cells, 6,424 cells (SSc) and 4,258 cells (Normal).(C-D) Images and quantificaton of Sun2 immunostaining in DAPI-stained skin sections of mice treated with PBS or bleomycin (Bleo) for 14 days. Scale bars = 100 *μ*m. (D) Histogram plot of Sun2 fluorescence in mouse skin treated with PBS or bleomycin (Bleo) for 14 days. N = 4 (PBS) or 3 (Bleo); n = 1271-4657 nuclei in each sample. Significance determined by Welch’s t-test. ****p < 0.0001.

To determine if the upregulation of *SUN2* represented a global increase in nuclear envelope components, we examined additional core components of LINC complexes—SUN2, LMNA, and SYNE1—to determine if the entire cytoskeletal to nuclear envelope bridge was upregulated in SSc patients. While SUN2 showed a profound and highly significant increase, its physical partners LMNA and SYNE1 remained statistically stable (Adjusted *p* > 0.05) (Fig.S1B-D). These data suggest that SUN2 upregulation is a specific, targeted transcriptional response in SSc fibroblasts, rather than a global up-regulation of transcripts encoding nuclear lamina components.

### SUN2 mediates nucleomechanosensing in dermal fibroblasts

To evaluate whether mechanical cues from the environment alter nuclear mechanosensing, we plated mouse dermal fibroblasts on collagen-coated polydimethylsiloxane (PDMS) substrates that were either soft (∼3 kPa) or stiff (∼1.5 MPa) for 24 hrs (Fig. 2A). Consistent with previous work ***Dupont et al. (2011***), YAP localized predominantly to the nucleus and the nuclear/cytoplasmic ratio of YAP fluorescence was significantly increased on stiff matrices compared with cells on soft substrates (Fig.S1A-B). Stiff substrates also elevated the expression of ECM and contractility genes, including *Col1a1, Col3a1*, and *Acta2*, and the matrix remodeling genes *Mmp8* and *Timp1*, relative to soft substrates (Fig.S1C).

**Figure 2.**
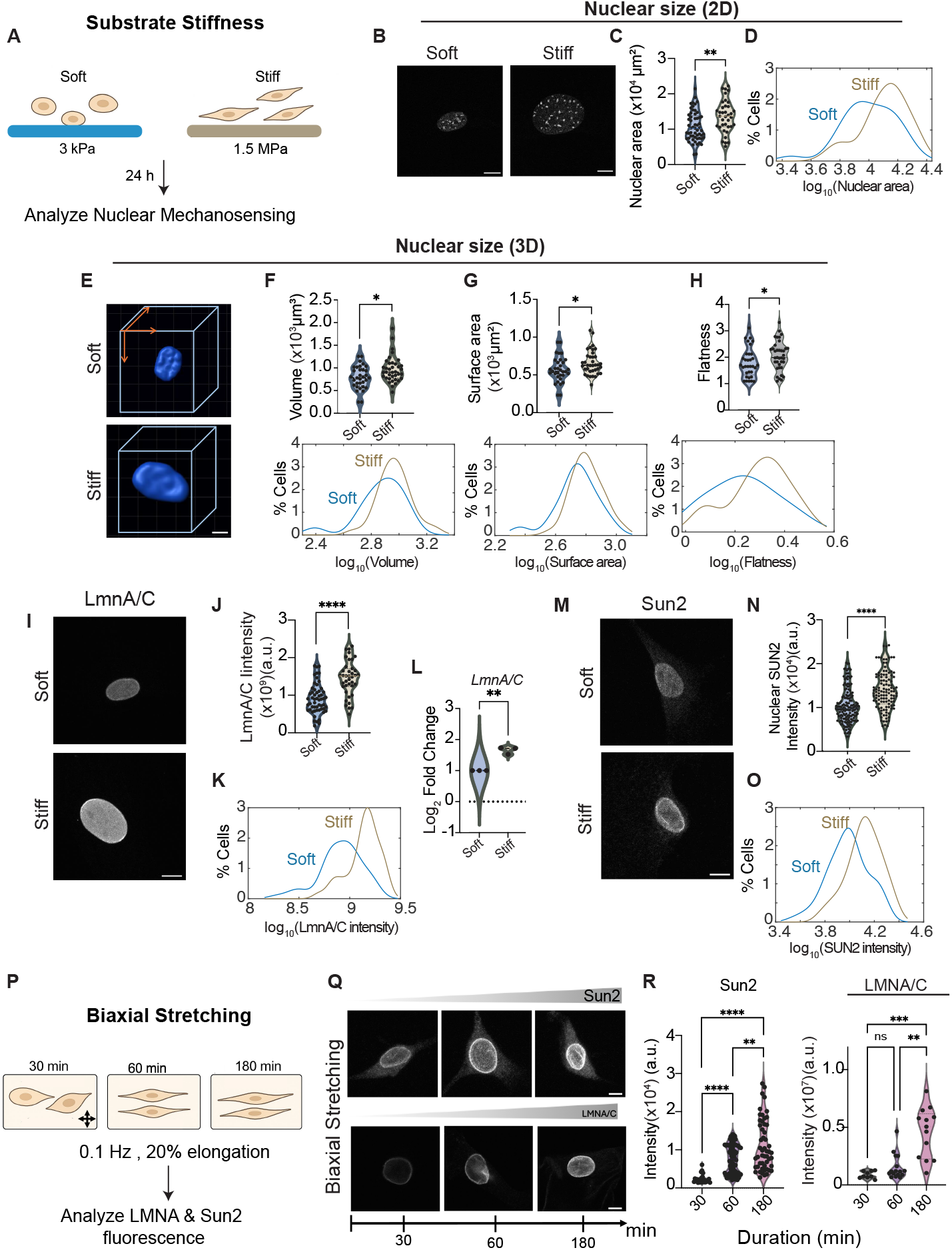
Nuclei respond to substrate stiffnessin dermal fibroblasts. (A) Schematic of an in vitro assay to examine nuclear mechanosensing in mouse fibroblasts. (B–D) Representative images and quantification of fibroblast DAPI-stained nuclei 2D size on soft or stiff substrates after 24 h. Scale bar = 10 *μ*m. Violin plot (C) and distribution (D) of fibroblast nuclear area on soft versus stiff substrates. *N* = 3, *n*_soft_ = 57, *n*_stiff_ = 37. (E-H) Representative images and quantification of 3D DAPI-stained nuclei on soft and stiff substrates. (F-H) Violin plots (top) and distributions (bottom) of nuclear 3D volume (F), surface area (G), and flatness (H) for mouse fibroblasts cultured on soft versus stiff substrates. *N* = 3, *n*_soft_ = 33, *n*_stiff_ = 37. (I-L) Representative images and quantification of A-type lamins (A-type lamins) in mouse dermal fibroblasts plated on soft and stiff substrates. *N* = 3, *n*_soft_ = 55, *n*_stiff_ = 39. (L) log_2_ fold-change in *LmnA* mRNA expression for mouse dermal fibroblasts on stiff versus soft substrates. (M-O) Representative images and quantification of Sun2 in mouse dermal fibroblasts plated on soft and stiff substrates. *N* = 2, *n*_soft_ = 169, *n*_stiff_ = 114. (P) Schematic of biaxial stretching experiments with mouse dermal fibroblasts. (Q–R) Images and quantification of A-type lamins and Sun2 in mouse dermal fibroblasts after biaxial stretching. Scale bar = 10 *μ*m. *N* = 2, *n*_t=30_ = 10 | 23, *n*_t=60_ = 16 | 72, *n*_t=180_ = 13 52 Significance determined by one way ANOVA. *p < 0.05, **p < 0.01, ***p < 0.001, ****p < 0.0001, ns = not significant.

We next asked how the stiffness of the substrate affects nuclear size, a key response to nuclear mechanosensing. The 2D area of dermal fibroblast nuclei was significantly elevated on stiff substrates compared with cells on soft substrates (Fig. 2B–D). 3D reconstructions further showed that stiff matrices increased nuclear volume and surface area and produced flatter nuclei, as reflected by higher flatness values and a rightward shift in the percentage cells cells with higher nuclear volume, surface area, and flatness (Fig. 2E–H). Sun2 and A-type lamins protein levels as well as *LmnA/C* transcripts were higher in mouse dermal fibroblasts on stiff compared to soft matrices (Fig. 2I-O). Sun2 also increased in human dermal fibroblasts in a mechanosensitive manner (Fig.S1D-E). By contrast, Sun1 levels were not elevated by stiffness in mouse fibroblasts (Fig.S1F-G). These data indicate that matrix stiffness is sufficient to remodel nuclear shape and increase a subset of nuclear lamina associated proteins in dermal fibroblasts.

To further test whether another model of mechanical stimuli, acute mechanical stretching, can modulate Sun2 and A-type lamins levels, we subjected mouse dermal fibroblasts to acute biaxial stretch (0.1 Hz, 20% elongation) (Fig. 2P) and monitored Sun2 and A-type lamin proteins over time.

Biaxial stretching induced a rapid and significant increase fluorescence intensity of both Sun2 and A-type lamins within 60 min, which was maintained or further enhanced at 180 min (Fig. 2Q–R). Thus, dynamic stretch increased Sun2 and lamin protein levels in fibroblasts, further showing Sun2 acts a mechanosensor in dermal fibroblasts.

### Loss of *Sun2* protects against fibrotic responses of skin fibroblasts in vivo and in vitro

To determine if *Sun2* is required for fibrosis progression in vivo, we treated control and *Sun2* KO mice with daily subcutaneous injections of bleomycin, which induces a robust fibrotic response in the skin ***Yamamoto et al. (1999); Utsunomiya et al. (2022); Caves et al. (2025***). Indeed, bleomycintreated WT mouse skin exhibited increased collagen remodeling and lipodystrophy compared to PBS treated mice at day 10 (D10) (Fig. 3A-C). By contrast, the skin of *Sun2* KO mice treated with bleomycin did not significantly increase collagen remodeling (Fig. 3A-C). At baseline, *Sun2* KO mice showed more perilipin fluorescence compared to WT mice (Fig. 3E; Fig.S2A). Following 10 days of bleomycin treatment, perilipin immunofluorescence in *Sun2* KO mouse skin showed a nonsignificant reduction compared to PBS-treated controls (Fig. 3D-E), suggesting that bleomycin may induce only modest lipodystrophy in the absence of *Sun2*, in contrast to the more almost complete lipodystrophy observed in WT mice. Thus, *Sun2* KO mouse skin exhibited a dampened response to bleomycin treatment compared to WT mice.

**Figure 3.**
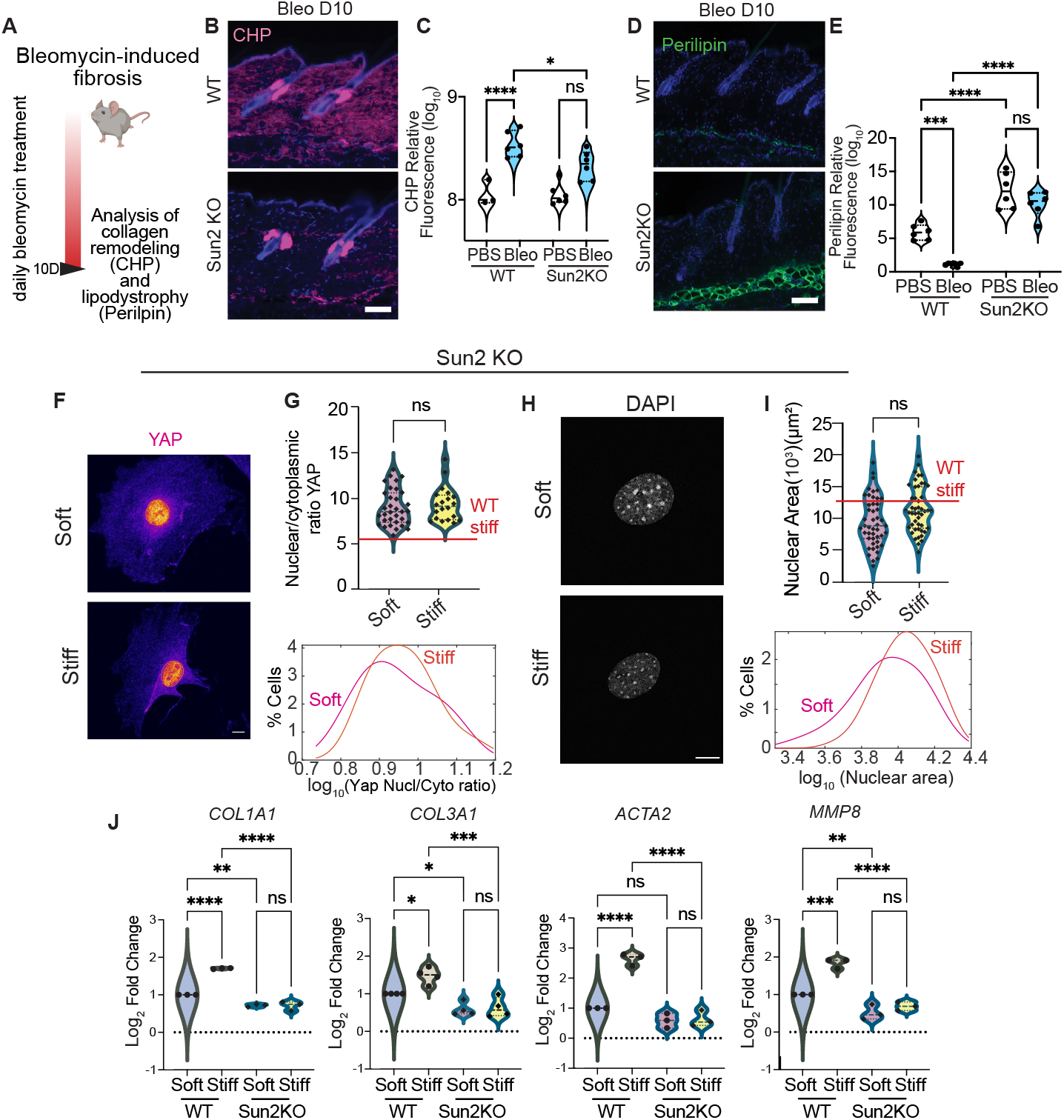
SUN2 is required for skin fibrosis in mice and fibroblast mechanosensing. (A) Schematic of bleomycin-induced skin fibrosis experiments in mice. (B-E)Representative images and quantification of skin sections from WT and *Sun2* KO mice treated bleomycin for 10 days stained with collagen hybridizing peptide (CHP, magenta) (blue) (A) or Perilipin antibodies (D, E) and DAPI. n= 5-6 mice. Significance determined by two way ANOVA. (F-G) Images and quantification of YAP localization in *Sun2* KO dermal fibroblasts plated for 24h on soft or stiff substrates. (F) Scale bars = 10. (G) Violin plot (top) and distribution (bottom) of the Yap nuclear/cytoplasmic ratio in SUN2 KO cells on soft versus stiff substrates (ns, not significant); the red line indicates the mean value for WT cells on stiff substrates. *N* = 3, *n*_soft_ = 33, *n*_stiff_ = 20. Significance was determined using a two-tailed Mann–Whitney U test. (H-I) Images and quantification of 2D nuclear size in *Sun2* KO dermal fibroblasts on soft and stiff substrates. (H) Representative DAPI stained fibroblasts. Scale bar= 10 *μ*m. (I) Violin plot (top) and distribution (bottom) of the 2D nuclear area of *Sun2* KO cells on soft versus stiff substrates; the red line shows the mean value for WT cells on stiff substrates. *N* = 3, *n*_soft_ = 47, *n*_stiff_ = 47. Significance was determined using a two-tailed Mann–Whitney U test. (J) Violin plots of log_2_ fold change in mRNA expression on stiff versus soft substrates for extracellular matrix genes (*Col1a1, Col3a1*), a myofibroblast contractility marker (*Acta2*), and a matrix-remodeling gene (*Mmp8*). N=3 independent experiments. Significance determined by one way ANOVA. *p < 0.05, **p < 0.01, ***p < 0.001, ****p < 0.0001, ns = not significant.

Next, we examined whether *Sun2* is required for fibroblast mechanosensing. As expected, WT fibroblasts exhibited stiffness-dependent increases in mean YAP nuclear translocation (Fig.S1A-B)and nuclear size (Fig. 2B). In contrast, *Sun2*-deficient fibroblasts cultured on soft versus stiff substrates showed similar mean YAP nuclear-to-cytoplasmic fluorescence ratios and comparable 2D nuclear areas (Fig.S2B-D). Notably, *Sun2* KO fibroblasts displayed nuclear YAP fluorescence (Fig. 3F) that was higher than WT cells on both soft and stiff substrates (Fig. 3G). Moreover, a greater fraction of *Sun2* KO fibroblasts showed a distribution of YAP nuclear-to-cytoplasmic ratios with high levels of nuclear YAP (Fig. 3G). These findings suggest that *Sun2*-deficient fibroblasts are mechanically activated even on soft substrates, consistent with the observation that lung fibroblasts and keratinocytes lacking *Sun2* are hypercontractile /cite Sandria***Stewart et al. (2015***) . Finally, the distribution of 2D nuclear areas revealed that more *Sun2* KO fibroblasts exhibited larger nuclei on stiff compared to soft substrates (Fig. 2B–D), indicating that *Sun2* KO fibroblasts retain the ability to respond to substrate stiffness.

The mean 3D nuclear volume, flatness, and surface area of *Sun2* KO fibroblasts were similar on soft and stiff substrates. However, the distribution of these parameters in *Sun2* KO fibroblast nuclei revealed a reduced nuclear surface area and 3D volume with no change in nuclear flatness on stiff substrates (Fig.S2B-D). Although the average A-type lamins fluorescence intensity did not differ between soft and stiff conditions in *Sun2* KO cells, a larger fraction of *Sun2* KO cells displayed higher A-type lamins fluorescence on stiff substrates compared to soft substrates (Fig.S2E-F).

Next, we examined whether stiffness-induced fibrotic gene expression depends on *Sun2*. In WT fibroblasts, fibrillar collagen transcripts (*Col1a1* and *Col3a1*), *Acta2* (encoding *α*-SMA), and the ECM-remodeling genes *Mmp8* and *Timp1* were upregulated on stiff compared to soft substrates (Fig. 3J and Fig.S2G). In contrast, *Sun2* KO fibroblasts exhibited lower expression of these transcripts under both soft and stiff conditions, with no detectable stiffness-dependent induction. Together, these findings indicate that stiffness-driven cellular responses including cytoskeletal contractility and nuclear size are translated via Sun2 to the chromatin for effective gene expression programs that drive ECM remodeling and fibrosis.

### *Sun2* is required for the transcriptional response to mechanical signals

To holistically examine how substrate stiffness reprograms dermal fibroblast transcription in a *Sun2*-dependent manner, we performed bulk RNA-sequencing on WT and *Sun2* KO skin fibroblasts cultured on soft or stiff substrates (Fig. 4A-D). Principal component analysis (PCA) of global gene expression revealed a strong separation between WT and *Sun2* KO fibroblasts along PC1, which accounted for 94% of the total variance, indicating that genotype is the primary driver of transcriptional differences. In contrast, samples cultured on soft versus stiff substrates showed a more modest divergence along PC2 (3% of variance). Biological replicates clustered tightly within each condition, supporting the reproducibility of the dataset (Fig. S3A).

**Figure 4.**
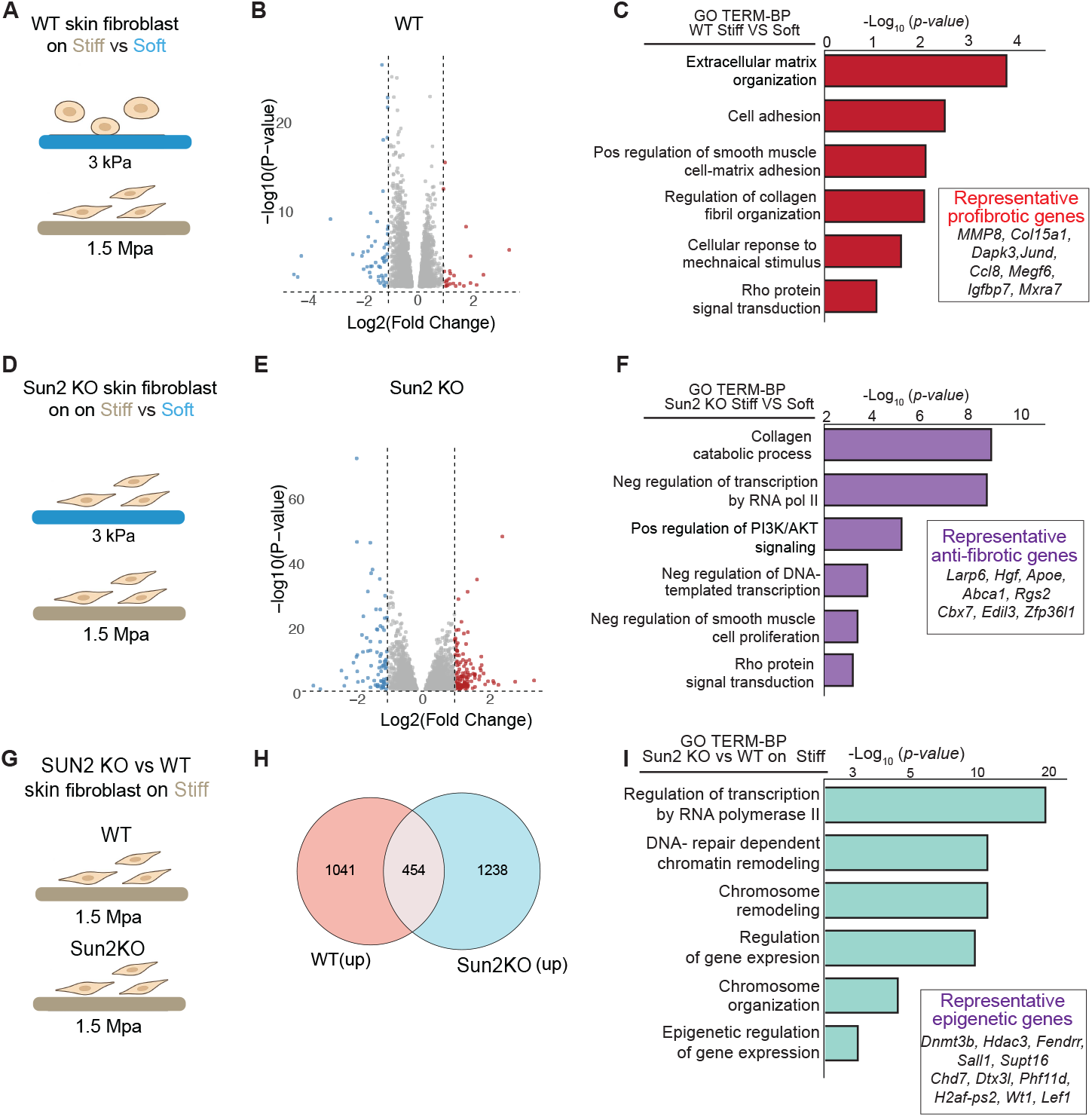
*Sun2* mediates epigenetic regulation of fibrotic gene expression. (A, D, and G) Schematic of in vitro stiffness experiment for RNA-seq: indicated genotype of dermal fibroblasts were cultured on soft (3 kPa) and/or stiff (1.5 MPa) collagen-coated substrates. (B and E) Volcano plots of differentially expressed genes in WT (B) or *Sun2* KO dermal fibroblasts on stiff versus soft substrates (log_2_ fold change vs log_10_ (p-value)); red, upregulated; blue, downregulated; grey, not significant. (C and F) Gene Ontology (GO) biological process (BP) enrichment plots for genes upregulated in WT (C) or or *Sun2* KO dermal fibroblasts (F) on stiff versus soft fibroblasts. (H) Venn diagram showing overlap between genes upregulated in WT fibroblasts on stiff versus soft substrates (WT up) and genes upregulated in *Sun2* KO versus WT fibroblasts on stiff substrates (*Sun2* KO up). (I) Gene Ontology biological process (GO TERM–BP) enrichment for genes upregulated in SUN2 KO versus WT fibroblasts on stiff substrates.

In WT fibroblasts on stiff substrates, 1495 genes were upregulated and 1503 genes were downregulated compared to fibroblasts on soft substrates (Fig. 4B). Analysis of mRNAs upregulated in WT cells on stiff substrates revealed using gene ontology: Biological Process (GO:BP) enrichment revealed induction of transcripts involved in extracellular matrix organization, cell adhesion, and collagen fibril organization (Fig. 4C). WT cells on stiff substrates downregulated transcripts involved in cell cycle, DNA repair, and chromatin organization (Fig. S3B). Gene expression changes in *Sun2* KO cells in response to stiff substrates were similar with 1692 transcripts upregulated and 1557 genes downregulated when plated on stiff substrates compared to soft substrates (Fig. 4E). Interestingly, transcripts upregulated in *Sun2* KO fibroblasts on stiff substrates were associated with collagen catabolic processing, negative regulation of transcription by RNA pol II, and positive regulation of PI3K/AKT signaling (Fig. 4F). *Sun2* KO fibroblasts downregulated genes involved in focal adhesion assembly, integrin signaling, and actomyosin organization (Fig. S3C).

Comparing the mRNAs upregulated in WT and *Sun2* KO fibroblasts on stiff substrates revealed a shared set of 454 mRNAs (Fig. 4H). mRNAs uniquely upregulated in *Sun2* KO fibroblasts on stiff substrates showed an enrichment in transcripts involved in epigenetic regulation of gene expression, chromosome organization, regulation of gene expression, and DNA repair–dependent chromatin remodeling (Fig. 4I).

### *Sun2* differentially regulates chromatin accessibility at mechanosensitive and profibrotic loci

Given that our data suggested a link between Sun2 and epigenetic remodeling during mechanosensing, we next asked whether *Sun2* regulates chromatin accessibility. We performed ATAC-seq on WT dermal fibroblasts cultured on soft and stiff substrates, as well as *Sun2* KO dermal fibroblasts cultured on stiff substrates (Fig. 5A). Analysis of differential peak locations between WT and *Sun2* KO fibroblasts on stiff substrates revealed a higher proportion of promoter-associated peaks and a lower proportion of intronic peaks in WT cells compared with *Sun2* KO cells (Fig. 5B). We compared mechanosensitive transcripts identified in WT dermal fibroblasts with ATAC-seq peaks enriched in WT cells relative to *Sun2* KO fibroblasts on stiff substrates. Peaks were annotated to promoters or to enhancers, defined here as intronic and distal intergenic regions excluding promoter-proximal loci. WT cells on stiff substrates exhibited significant enrichment of accessible peaks at both promoters and enhancers associated with mechanosensitive genes, with enhancer–transcript overlap showing greater statistical significance than promoter–transcript overlap. These data indicate that *Sun2* loss is associated with altered chromatin accessibility in response to mechanical stimulation.

**Figure 5.**
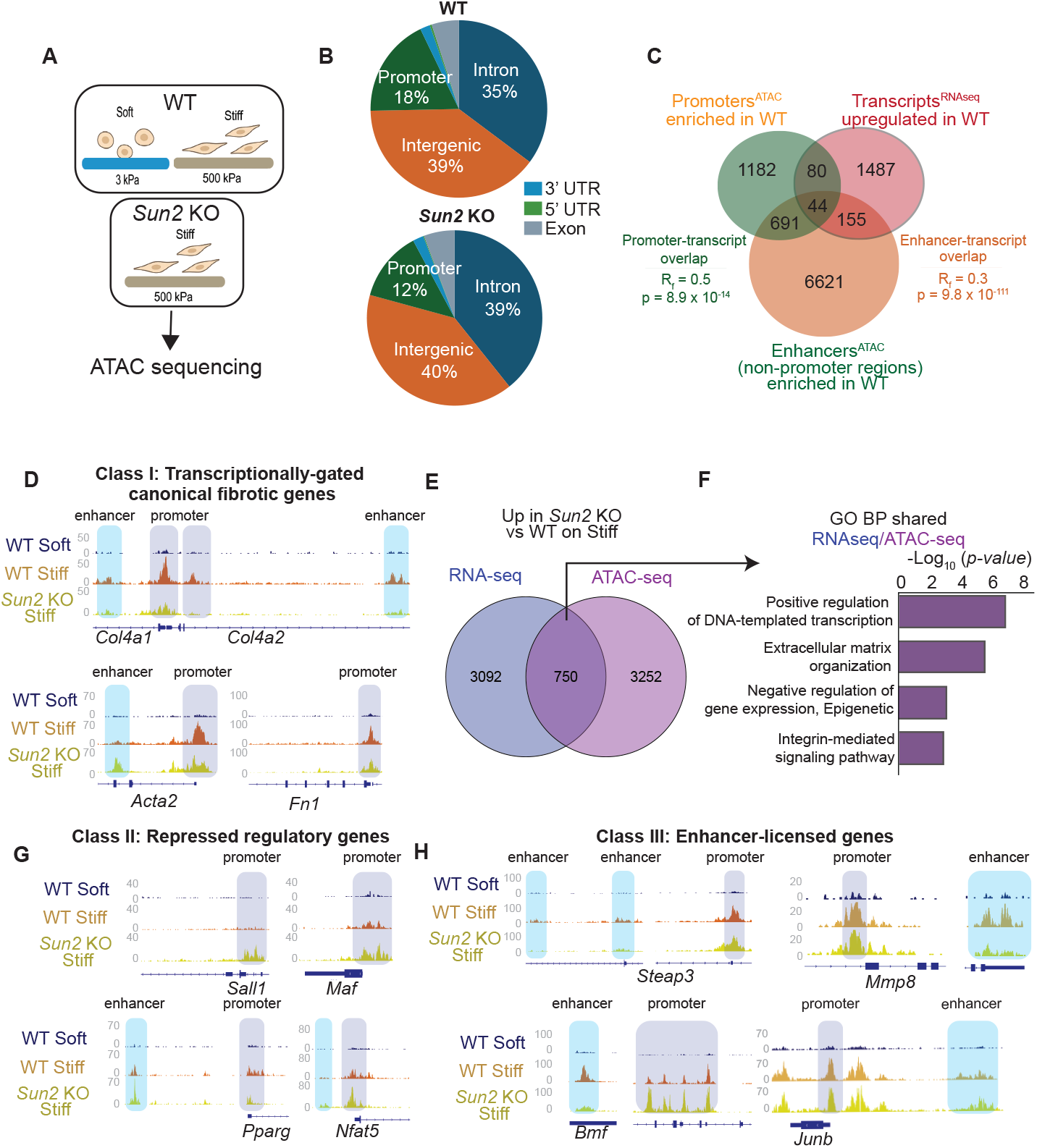
*Sun2* mediates epigenetic changes in fibrotic gene expression. (A) Schematic of ATAC sequencing experimental design. (B) Pie chart of annotated genomic regions enriched in 10,647 loci in WT dermal fibroblasts and 8243 loci in *Sun2* KO cells. (C) Venn diagram illustrating the overlap of mRNAs upregulated in WT dermal fibroblasts on stiff vs soft substrates (TranscriptsRNAseq) (see Figure 4B) with enhancers and promoters enriched in WT vs Sun2 KO cells identified in ATAC-seq data. Enhancers were defined as intronic and distal intergenic ATAC-seq peaks. Statistical analysis performed using hypergeometric test. (D)Snapshots of IGV tracks from ATAC sequencing data for canoncial fibrotic genes. (E) Venn diagram illustrating overlap between mRNAs upregulated in *Sun2* KO versus WT fibroblasts on stiff substrates (RNA-seq; blue) and loci with increased chromatin accessibility (ATAC-seq; purple). (F) Gene Ontology (GO) biological process analysis of genes upregulated in *Sun2* KO fibroblasts and loci with increased accessibility compared to WT cells on stiff substrates. (G) Screenshots of IGV tracks from ATAC sequencing data for transcriptional regulatory genes upregulated in *Sun2* KO cells compared to WT dermal fibroblasts. (H) Snapshots of IGV tracks from ATAC sequencing data for mechanosensitive genes in WT dermal fibroblasts.

We first examined genes canonically implicated in fibrosis, including *Col* genes, *Acta2*, and *Fn1*, which are not induced by stiff substrates in *Sun2* KO fibroblasts (Fig. 4J). Despite the lack of transcriptional induction, *Sun2* KO fibroblasts cultured on stiff substrates displayed differential chromatin accessibility of promoters and, in some cases, distal regulatory regions of these genes (Fig. 5C). These findings suggest that canonical fibrotic genes maintain accessible chromatin but require *Sun2* for transcriptional activation in response to mechanical cues.

To further assess the relationship between chromatin accessibility and gene expression, we identified 750 genes that were transcriptionally upregulated and exhibited increased chromatin accessibility in *Sun2* KO fibroblasts on stiff substrates. These genes were enriched for transcriptional and epigenetic regulators, as well as ECM and integrin remodeling genes (Fig. 5E). Regulatory genes that are anti-fibrotic such as *Gli2* ***Yan et al. (2021); Kramann et al. (2015***), *Pparg* ***Feng et al. (2022); Zhang et al. (2023***), and *Sall1* displayed increased promoter and enhancer accessibility in *Sun2* KO fibroblasts on stiff substrates but not in WT fibroblasts under the same conditions (Fig. 5F), indicating that Sun2 normally restrains the accessibility of the promoters of these anti-fibrotic regulatory loci.

Finally, we examined loci for genes induced by stiff substrates in WT dermal fibroblasts with lower peaks in enhancer regions in *Sun2* KO fibroblasts (Fig. 4C), indicating that these Sun2 regulates cis-regulatory loci of these genes. Both WT and *Sun2* KO fibroblasts exhibited promoter accessibility at several fibrotic gene loci under stiff conditions including *Junb* ***Pi et al. (2022***), *Mmp8* ***Salo et al. (2014); Naim and Baig (2020); García-Prieto et al. (2011***), and *Steap3* ***Li et al. (2020); Wang et al. (2025***); however, distal enhancer accessibility was selectively observed in WT fibroblasts and was markedly reduced in *Sun2* KO fibroblasts (Fig. 5G). These data indicate that Sun2 is required for enhancer activation at a subset of mechanosensitive genes.

Collectively, these results define three mechanoregulated chromatin states regulated by Sun2: (1) canonical fibrotic genes that require SUN2 for transcriptional activation despite accessible chromatin, (2) transcriptional and epigenetic regulatory genes that become derepressed upon *Sun2* loss, and (3) mechanosensitive genes whose activation depends on SUN2-dependent enhancer accessibility.

### Sun2-Ezh2 axis regulates fibrotic gene expression

Based on these findings and previous studies implicating Ezh2 in the regulation of fibrotic genes in fibroblasts ***Le et al. (2021); Tsou et al. (2019***), we next investigated whether SUN2 regulates Ezh2 expression in response to matrix stiffness. Immunofluorescence analysis revealed that WT fibroblasts exhibited increased Ezh2 protein levels when cultured on stiff compared with soft substrates. In contrast, *Sun2* KO fibroblasts failed to upregulate Ezh2 in response to stiffness and instead displayed a pronounced reduction in Ezh2 protein abundance (Fig. 6A–B). Consistent with these findings, qPCR analysis demonstrated increased expression of *Ezh2* in WT fibroblasts on stiff matrices, while *Sun2* KO fibroblasts did not show stiffness-dependent induction of *Ezh2* transcripts (Fig. 6C). Thus, Sun2 is required for stiffness-dependent induction of Ezh2 expression in mouse dermal fibroblasts.

**Figure 6.**
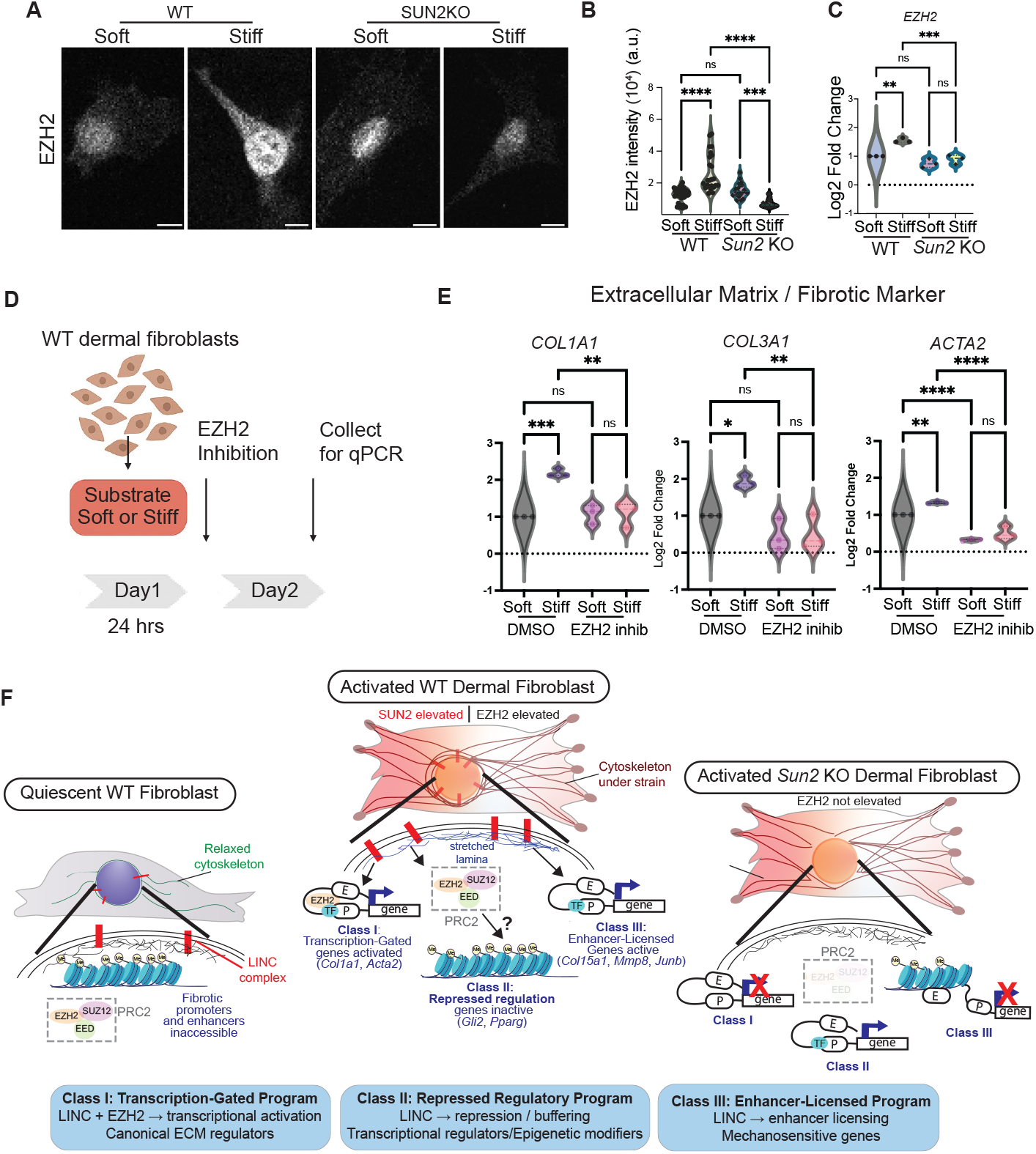
*Sun2* Ezh2 axis regulates fibrotic gene expression. (A-C) Images and quantification of Ezh2 immunofluorescence in WT and *Sun2* KO dermal fibroblasts cultured on soft or stiff substrates. (A) Scale bar = 10 *μ*m. (B) Quantification of nuclear Ezh2 intensity. *N* = 2, *n*_soft_ = 21–37, *n*_stiff_ = 20–24. Significance was assessed using a two-tailed Mann–Whitney U test. (C) qPCR analysis of *Ezh2* mRNA expression in WT or *Sun2* KO fibroblasts on soft or stiff substrates. *N* = 3. Significance was assessed using a two-tailed Mann–Whitney U test. (D-E) Schematic and analysis of WT and *Sun2* KO dermal fibroblasts plated on soft or stiff substrates for 24 h, followed by 24 h treatment with the Ezh2 inhibitor GSK343 or DMSO (Day 1), and collection for qPCR (Day 2). *N* = 3 independent experiments. Statistical significance was assessed using one-way ANOVA. ns, not significant; *p* < 0.05, \**p* < 0.01, \*\**p* < 0.001, \*\*\**p* < 0.0001. (F) Proposed model for SUN2–LINC–PRC2–dependent control of profibrotic gene expression in dermal fibroblasts. In quiescent WT fibroblasts, fibrotic genes are repressed. Upon mechanical stimulation of WT cells, cytoskeletal tension transmitted through LINC complexes promotes lamin stretching and altered Ezh2-driven epigenetic remodeling to activate transcription, repress transcriptional activators, and promote enhancer-licensing. In *Sun2* KO dermal fibroblasts,loss of LINC complexes blunts this mechanotransduction despite cytoskeletal activation.

To determine the functional contribution of Ezh2 to fibroblast mechanosensing, we pharmacologically inhibited Ezh2 in dermal fibroblasts cultured on soft and stiff matrices and assessed fibrotic gene expression by qPCR (Fig. 6D). Ezh2 inhibition significantly attenuated the stiffnessinduced upregulation of *Col1a1, Col3a1*, and *Acta2* in fibroblasts cultured on stiff substrates (Fig. 6E). Together, these data support a model in which Sun2-dependent chromatin accessibility changes engage an Ezh2-mediated epigenetic program that is required for profibrotic gene expression for canonical fibrotic genes in response to increased matrix stiffness (Fig. 6F).

## Discussion

Our data indicates that mechanosensing in dermal fibroblasts requires the integral inner nuclear membrane protein Sun2, which comprises the nuclear aspect of a subset of LINC complexes (Fig. 6F). One mechanism by which Sun2 can act in the response of fibroblasts in fibrosis is through its interactions with A-type lamins, which increases in dermal fibroblasts on stiff substrates. The expression of A-type lamins is well known to scale with matrix stiffness ***Swift et al. (2013***), and Sun2 via LINC complexes can directly bind to A-type lamins within the nucleus ***Chang et al. (2015b***). We previously demonstrated that integrin-mediated adhesions transmit high mechanical tension to LINC complexes and A-type lamins in the nuclear lamina ***Carley et al. (2021***). The current data resonate with the ability of cyclic stretch to upregulate lamin A in lung fibroblasts in vivo during mechanical changes associated with ventilation ***López-Alonso et al. (2018***). Finally, mechanosensitive changes in fibroblasts via lamins have been linked to chromatin remodeling and changes in gene expression in fibroblasts ***López-Alonso et al. (2018***).

We propose that fibrotic cues upregulate SUN2, remodel the nuclear lamina, and alter the chromatin and transcriptional machinery to drive gene expression changes during fibrosis. These findings resonate with our prior work demonstrating that SUN proteins suppress chromatin remodeling in keratinocytes during differentiation ***Carley et al. (2021***), and as well as studies showing that Nesprin-2–deficient dermal fibroblasts show reduced H3K9me3 ***Amiad Pavlov et al. (2023***). Tension on Sun2 may change the diameter of nuclear pore complexes, which shows some LINC-complex dependent dilation ***Morgan et al. (2025***) to control mechanosensitive transcriptional factor translocation ***Elosegui-Artola et al. (2017***). However, our data indicate that nuclear localization of the mechanosensitive transcription factor YAP is slightly elevated by *Sun2* loss in fibroblasts on stiff substrates. In parallel, Sun2 could also regulate gene accessibility, silencing, and activation directly in fibrosis given that Sun proteins are directly tethered to chromatin ***Amiad Pavlov et al. (2023); Chen et al. (2018); Lityagina and Dobreva (2021***). In this context, Sun2 might modulate chromatin–lamina interactions, positioning pro-fibrotic loci for regulation by mechanical cues and associated transcription factors.

This work further implicates Ezh2 as a key effector of fibrosis and indicates that Sun2 regulates Ezh2 in a mechanosensitive-chromatin axis. Ezh2 has traditionally been characterized as a transcriptional repressor that acts through deposition of H3K27me3; however, emerging evidence indicates that Ezh2 also plays context-dependent roles in transcriptional activation in breast and prostate cancer***Kim et al. (2018); Gonzalez et al. (2011); Anwar et al. (2023***). Our data support a model in which Ezh2 acts downstream of Sun2 to enable fibrotic gene expression in response to increased matrix stiffness, rather than simply enforcing chromatin repression. How might Sun2 regulate Ezh2 and its targets? One possible mechanism is through Sun2-lamin interactions which can alter Ezh2 activity or its chromatin interactions. For instance, in muscle, loss of A-type lamins can promote disassembly of polycomb protein foci and protein dispersion ***Cesarini et al. (2015***) and increase the accessibility of Ezh2 chromatin, mainly in promoter regions ***Jabre et al. (2025***). Future work will investigate whether SUN2 directly modulates Ezh2-chromatin interactions during fibrotic gene expression.

An alternative, not mutually exclusive, mechanism is that Sun2 influences nuclear–cytoskeletal coupling and intracellular signaling pathways that regulate Ezh2 expression and activity. For example, changes in force transmission through LINC complexes could change *PI3K/AKT* expression or activity, which we found was dysregulated in *Sun2* KO cardiac cells ***Stewart et al. (2019***). This pathway and its related mechanosensitive signaling that has been linked to Ezh2 stability and function ***Riquelme et al. (2016***), thus could be contributing to the altered transcriptional state in *Sun2* KO fibroblasts in response to stiff substrates. Consistent with this idea, our ATAC-seq and RNA-seq data show an enrichment for PI3K/AKT signaling and transcriptional regulators in *Sun2*-deficient cells, suggesting a rewiring of mechanosensitive signaling downstream of *Sun2* loss.

Given the central role of dermal fibroblasts in cutaneous fibrosis and systemic sclerosis, our prior work implicating *Sun2* in cardiac fibrosis ***Stewart et al. (2015***) and a companion article on its role in lung fibrosis, we propose that suppressing nuclear mechanotransduction, particularly the Sun2–Ezh2 mechano-epigenetic axis, may represent a promising therapeutic node for multi-organ fibrotic therapies. Targeting SUN2 itself, or modulating downstream epigenetic regulators such as Ezh2, could potentially attenuate pathological matrix production while preserving the necessary reparative responses. Future studies defining cell type–specific functions of Sun2, defining specific epigenetic post-transcriptional modifications in response to mechanical stimuli, and testing whether similar mechano-epigenetic coupling operates in other fibrotic tissues will be important for understanding how broadly this mechanism contributes to fibrosis and how best to target it therapeutically.

## Methods

### Sex as a biological variable

Our study exclusively examined male mice. It is unknown whether the findings are relevant for female mice.

### Animals

Male, wild-type C57BL6/J mice were purchased from Charles River (stock 027) and *Sun2* KO mice (Jax stock 012716) mice were purchased from The Jackson Laboratories. To avoid variations in DWAT associated with the hair cycle, male mice were used to avoid variable hair cycling in female mice, and we confirmed that mice were in the 2^nd^ telogen as indicated by a lack of follicular pigment in the backskin. Animals were maintained on a standard chow diet (Harlan Laboratories, 2018S) in a 12-hour light/dark cycle. All animal care and experiments followed guidelines issued by Yale University’s Institutional Animal Care and Use Committee (IACUC).

### Chemical Induction of Fibrosis

Male mice between 6-8 weeks received daily subcutaneous injections of 300ug Bleomycin Sulfate (Enzo) for indicated time frame as described in ***Caves et al. (2025***).

### Generation of hydrogels

Hydrogels were prepared by mixing base and cross-linker components (CY 52-276A/B for soft gels; Sylgard 184, Dow Corning for stiff gels). Soft (3 kPa) gels were mixed at a 1:1 ratio and stiff (1.5 MPa) gels at a 10:1 ratio. Mixtures were immediately degassed for 15 min. Surfaces of 35 mm culture dishes (MatTek Life Sciences) were covered with 200 µL of the desired gel and spin-coated at 1000 rpm for 60 s (Headway Research, PWM32). Substrates were cured at 60°C overnight (soft) or for 1 h (stiff). Prior to use, substrates were sterilized under UV light for 30 min and treated with a plasma cleaner for 1 min. Collagen was incubated on the hydrogels for 1 hour at 37°C. Dermal fibroblasts were plated overnight, and then fixed for 10 min in 4% paraformaldehyde (Merck).

### Stretching

For biaxial stretch, culture plates with a silicon elastomer membrane (Flexcell, BF-3001U) were coated with collagen (50 micrograms/ml) in PBS for 1h at 37°C prior to cell seeding. 60 000 cells per elastomer were seeded 24h prior to experiment start. Cells were then exposed to cyclic mechanical strain using the Flexcell Tension System (FX-6000T_TM_; FlexCell International Corporation) at 20% elongation, 0.1Hz frequency. Stretching was applied at 37°C, 5% CO2 for 30, 60 and 180 min.

### Immunofluorescence staining

Mouse back skin from chemically induced fibrosis was embedded in O.C.T. and sectioned across the hair follicle. 14mm cryosections were collected and stained with H&E or trichrome as described by Polysciences Inc. For immunofluorescence staining, 14mm cryosections were processed as previously described in ***Caves et al. (2025***). Briefly, slides were fixed for 10 minutes in 4% PFA, washed and blocked in 10% normal donkey serum (Jackson Immunology) for 1 hour at room temperature. Slides were stained for Plin (Abcam, ab61682) overnight at 4°C.

Mouse dermal fibroblasts were fixed with 4% paraformaledyde (PFA) for 10 minutes at room temperature, followed by three 5 minute PBS wash cycles. Samples were subsequently blocked for 1 hour at room temperature with a blocking solution consisting of 1 mL normal donkey serum (NDS, Jackson Immuno, 017-000-121) and 400 µL 10% Triton-X in 8.6 mL PBS. Primary antibody diluted in blocking solution – SUN2, 1:200 (Abcam, AB124916), YAP, 1:200 (CST, 4912S), LaminA/C (Abcam, AB133256), and/or Ezh2, 1:200 (CST, 5246T) was incubated overnight at 4°C. Primary incubation was followed by one minute PBS wash cycles at room temperature. Primary antibody solution was replaced with secondary antibody solution diluted – 1:200 to 1:1000 – in blocking solution, in which samples were immersed for 1 hour at room temperature, followed by one minute PBS wash at room temperature. Nuclei were labeled with HOECHST 34745.

### Collagen hybridizing peptide

Hybridization with collagen hybridizing peptide was performed as described ***Caves et al. (2025***). Cryosections (14 µm) were allowed to equilibrate to room temperature prior to vigorous washing with PBS to remove residual OCT embedding medium. Sections were subsequently blocked with 5

### Imaging

For H&E, tissue sections were imaged at 10x using a Zeiss AxioImager M1 (Zeiss) equipped with a Zeiss AxioCam MRc. CHP fluorescence was imaged at 40x using a Zeiss AxioImager M1 (Zeiss) equipped with an Orca camera (Hamamatsu). B-CHP images were analyzed as described in ***Caves et al. (2025***).

Fibroblasts in vitro were imaged with a Zeiss LSM 880, a confocal laser scanning microscope at the Light Microscope Imaging Facility on Science Hill, Yale University. A 40×, 1.45 NA objective was used for imaging. 3D stacks of images were acquired with 250 nm slice thickness. For multi-channel imaging, pinhole diameter was adjusted to achieve equal slice height across channels. Nuclei were imaged in the DAPI channel (405 nm) channel; actin cytoskeleton was imaged in the GFP channel (488 nm) and the immunofluorescence-stained sample was imaged in the Cy5 (642 nm) channel.

### RT-qPCR

Total RNA was isolated using the RNeasy Plus Micro Kit (Qiagen). RNA concentration was measured using a Nanodrop™One Spectrophotometer (ThermoFisher Scientific). cDNA was transcribed from equal amounts of RNA (500 ng) using the SuperScript™III First-Strand Synthesis System (ThermoFisher Scientific). Amplification reactions were performed with the LightCycler 480 SYBR Green I Master Mix on the corresponding LightCycler 480 II system (Roche Diagnostics). All reactions were conducted in triplicate with the following program: 95°C for 10min, followed by 45 cycles of 95°C for 10s, 60°C for 10s, and 72°C for 10s. GAPDH was used to normalize expression levels of the genes of interest. Primer sequences were generated by the Harvard PrimerBank and synthesized by Eurofins. Relative gene expression in data was quantified using the 2^−ΔΔ*CT*^ method.

**Table 1.**
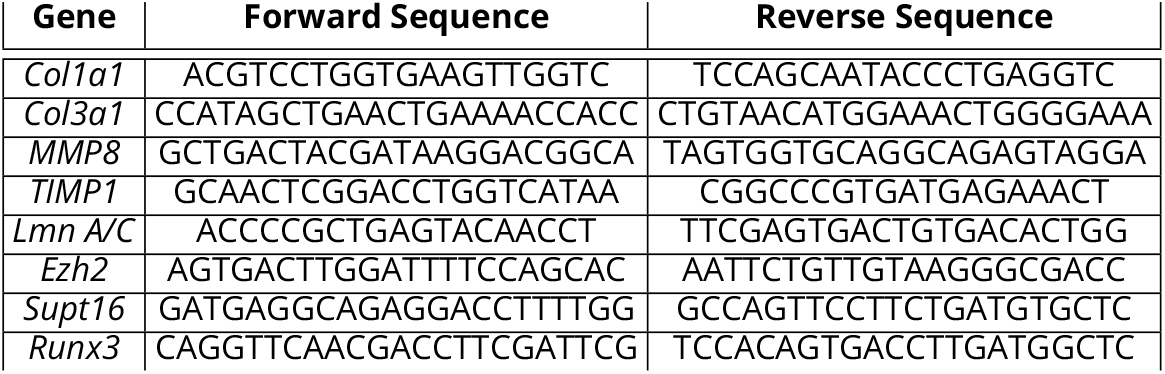
Mouse RT-qPCR Primers.

### Image Analysis

Microsopic images were analyzed using MATLAB R2024b. First, maximum intensity projections were calculated from Z-stacks, resulting in a single image for each channel. Then, nucleus images (DAPI channel) were Gaussian-smoothed with *σ* = 5 pixels and then binarized using the *imbinarize* routine with an appropriate threshold (e.g., 2× or 5× median pixel) and a minimum area threshold. Thresholds were adjusted based on manual inspection of the segmented nuclei. For each segmented nucleus, various parameters such as area, perimeter, and circularity, were calculated using the *regionprops* function. When needed, cells were similarly segmented from the actin channel. These segmentations were used to quantify the intensity within or outside nucleus, for any channel. Median intensity was reported as the Intensity for each channel, after subtracting the median intensity of the background (pixels not segmented as nucleus or cell). Nuclear aspect ratio was calculated as the ratio of the long axis to short axis, after obtaining ellipse fits for the nucleus boundaries. For 3D analysis, we used NucleusJ 2.0, an automated ImageJ plugin for 3D bio-imaging analysis ***Dubos et al. (2020***).

### Fibroblast Isolation

Primary human dermal fibroblasts were isolated from skin tissue using an enzymatic dissociation workflow. Fresh tissue was transported on ice in DMEM/F12 supplemented with 1% antibiotic antimycotic, washed briefly in 10% Betadyne and then washed three times in culture medium (1 min each), and minced into ∼0.2 cm^2^ pieces. Minced tissue was digested in DMEM (high glucose) containing Liberase, collagenase I, collagenase II, DNase, HEPES (25 mM), sodium pyruvate, non-essential amino acids, and penicillin/streptomycin/amphotericin. Digestion was performed at 37°C for 1 h with agitation. The suspension was filtered through a 70 µm mesh, washed with PBS, and pelleted by centrifugation (300×*g*, 10 min, 4°C). When excessive red blood cells were present, pellets were treated with sterile ACK lysis buffer on ice for 1 min, diluted with ice-cold PBS, and repelleted. Cells were resuspended in DMEM containing 10% FBS and 1% antibiotics, counted, and plated on adherent culture dishes.

### Human scRNA-sequencing

Skin biopsies from 5 patients with scleroderma (SSc1-5) and 6 healthy controls (Normal1-6) were analyzed using 10x genomics platform. Patient selection and criteria, as well as sample processing was described in Odell et al. and written, informed consent was obtained (Odell et al., 2022). Briefly, 6mm biopsies were collected and incubated in RPMI 1640 medium (Gibco) containing 5% fetal bovine serum (5% FBS/RPMI) and 10 mg/ml Dispase II (Sigma D4693–1G) for 45 minutes at 37° C shaking at 200–250 rpm. The 6 mm sample was then removed from the media and minced with sterile iris scissors, followed by digestion with Liberase TM (Sigma) 0.5 mg/ml and DNase I 30 Units/ml in 5% FBS/RPMI for 45 minutes at 37° C shaking at 200–250 rpm. The resulting single cell suspension was filtered through a 70 m nylon membrane and washed. Live cells were sorted on a FACSAria at the Yale Flow Core and their final concentration and viability quantified with Trypan blue on a hemacytomer. The cells were pelleted and suspended in phosphate buffered saline containing 0.04% bovine serum albumin between 500–1000 cells/l. 3000–6000 cells with greater than 80% viability were submitted to the Yale DNA Sequencing facility for generation of single-cell cDNA libraries using the Chromium Single Cell Controller (10x Genomics). The scRNA-seq processing was described in O’Dell et. al. (Odell et al., 2022).

### RNA seq

Six million cells were plated onto the desired stiffness substrate 24 hours prior to conducting the RNA sequencing experiment. Samples were digested using Trizol LS (Invitrogen). After isolation, lysates were immediately placed on ice and stored at -80C until RNA purification. RNA purification was carried out using the RNeasy Plus Micro Kit (Qiagen) following manufacturer instructions. RNA concentrations were assessed using Qubit fluorometer (Q33238; Thermo Fisher Scientific). All samples were sequenced by the Illumina Next-seq 550 instrument with paired-end 2×75bp read length and sequenced to a depth of at least 25M reads per sample in the Yale Center for genome analysis.

Bulk RNA-seq data were processed using a standard alignment-based workflow. Following initial quality control, adapter sequences were trimmed from raw reads prior to alignment. Reads were aligned to the reference genome using STAR, and gene-level read counts were generated with featureCounts by assigning reads to annotated exons. For downstream analyses, genes were filtered by excluding any gene for which at least one sample had fewer than 50 counts; count-distribution plots were generated before and after filtering to assess the impact of this threshold. Sample-to-sample relationships were evaluated by principal component analysis (PCA) using the first two principal components. Differential expression analysis was performed in R using DESeq2, and results were reported for the *Sun2* KO versus WT contrast and soft vs stiff substrates. Functional interpretation of differentially expressed genes was performed using over-representation analysis (ORA) and gene set enrichment analysis (GSEA) implemented in clusterProfiler, with pathway visualization performed using pathview.

### Ezh2 Inhibition

Fibroblasts were seeded on soft and stiff hydrogels at equal concentrations (1,000,000 cells/mL). Twenty-four hours post-seeding, Ezh2 was inhibited using GSK343 (Sigma, SML0766) at 10 *μ*m. RNA samples were then collected 24 hours after inhibition and gene expression levels were analyzed using RT-qPCR.

### ATAC seq

ATAC-sequencing libraries were generated as previously described ***Bao et al. (2015***). Briefly, 50,000 murine dermal-derived fibroblasts were seeded onto 6 cm dishes coated with 3 or 1.5 MPa PDMS substrates and incubated for 24 h prior to tagmentation. Cells were washed twice with PBS and lysed in resuspension buffer (10 mM Tris-HCl, pH 7.5, 10 mM NaCl, 3 mM MgCl_2_) supplemented with 0.1% NP-40, 0.1% Tween-20, and 0.01% digitonin for 10 min on ice, as previously described ***Buenrostro et al. (2015***). Tagmentation buffer containing Illumina Tn5 transposase (25 *μ*L TD buffer; Nextera Kit, Illumina, San Diego, CA, USA), 5 *μ*L TDE1 transposase, and 20 *μ*L nuclease-free water was added to fibroblasts adhered to the culture dish. Samples were incubated for 30 min at 37 ^◦^C on a shaker. DNA was purified using the Qiagen MinElute Reaction Cleanup Kit according to the Kaestner Lab protocol. Library fragment size distributions were assessed using an Agilent Bioanalyzer through the Yale Keck Biotechnology Resource Laboratory, and samples were submitted to the Yale Center for Genomic Analysis for sequencing. Promoter and enhancer loci were validated using annotations from Ensembl (release 114)(https://ensembl.org).

### Processing of samples

Sequencing reads were quality-checked using FastQC and aligned to the mouse reference genome (mm39) using Bowtie2. Aligned reads were filtered to remove low-quality reads (MAPQ < 30), duplicate reads, and mitochondrial reads. Fragment size distributions and transcription start site (TSS) enrichment scores were calculated as quality control metrics. Peaks representing regions of accessible chromatin were identified using MACS2. Peaks were visualized in the Integrated Genome Viewer (IGVWeb) ***Robinson et al. (2011***).

### Analysis of Single Cell RNA sequencing data

For human SSc samples (GEO: GSE214088), we performed an unbiased screen of baseline fibrotic markers. Three control samples (Normal 2, 4, and 5) exhibited *COL1A1* levels exceeding the median of the SSc cohort, suggesting underlying pathology or localized scarring. To prevent these ‘activated’ controls from masking disease-specific signatures, we used healthy 1 and 3 for our primary analysis of mRNA differences in fibroblasts.

### Statistics

Data are presented as mean ± SD or as violin plots with lines at the median (center) and 25/75% quartile (top and bottom). Group sizes were determined based on the results of preliminary experiments and mice were assigned at random to groups. Experiments were not blinded. Statistical significance for each experiment was determined as shown in the figure legends, where N = the number of individual experiments, n = the number of independent biological replicates (animals or cells) per group. Data between two groups were analyzed using a two-sided Mann–Whitney U test. Data between more than two groups were statistically compared using a one-way or two-way analysis of variance (ANOVA), as appropriate. RT–qPCR measurements were obtained from three independent experiments (*N* = 3). Within each experiment, each target was measured in technical triplicate and averaged; the averaged value was used for downstream analysis and plotting. Signficance of gene overlap determined by hypergeometric test. Statistical significance of bulk RNA-seq data was calculated using an adjusted p-value cutoff <0.05 in the DESeq2 R package. Statistical analyses were performed in Prism (GraphPad), DESeq2, or R.

## Data availability

Genomic data have been deposited at GEO as GSE315855 and GSE315855. All data are publicly available as of the date of publication. This paper also analyzes existing, publicly available data, accessible at GEO: GSE214088. Other data reported in this paper will be shared by the lead contact upon request.

## Study approval

All animal care and experiments were approved by Yale University’s Institutional Animal Care and Use Committee (IACUC).

## Author contributions

Conceptualization, A.N., S.S., K.D., S.D., M.K., M.H., and V.H.; Methodology, A.N., K.D., S.S.; Investigation, A.N., K.D., S.S., S.D., and V.A.T.; Formal analysis, A.N., J.D.A, R.R., Visualization: A.N., S.S., K.D., and V.H.; writing—original draft, A.N. and V.H.; writing—review & editing, A.N., K.D., S.S., J.D.A., M.K., and V.H.; funding acquisition, M.K. and V.H.; supervision, M.K. and V.H.

## Funding support

Yale Center for Genomic Analysis receives funding from NIH Shared Equipment grant #1S10OD028669-01. S.S is funded by 1F31 AR085488. V.H. is funded by NIH NIAMS R01s AR076938, AR0695505, and AR084558, the Leo Foundation, and Boehringer-Ingelheim Pharmaceuticals, Inc. M.K is funded by R35 GM153474 and Boehringer-Ingelheim Pharmaceuticals, Inc.

## Acknowledgments

We thank the Yale Science Hill Imaging Facility and the Yale Center for Genomic Analysis for support and assistance in this work. We thank Axel Poulet for 3D image analysis assistance.

## Notes

### Competing Interest Statement

The authors receive research funding from Boehringer-Ingelheim Pharmaceutical, Inc.

