## Supplemental Figures for "SUN2 mediates epigenetic remodeling to drive mechanotransduction during skin fibrosis"

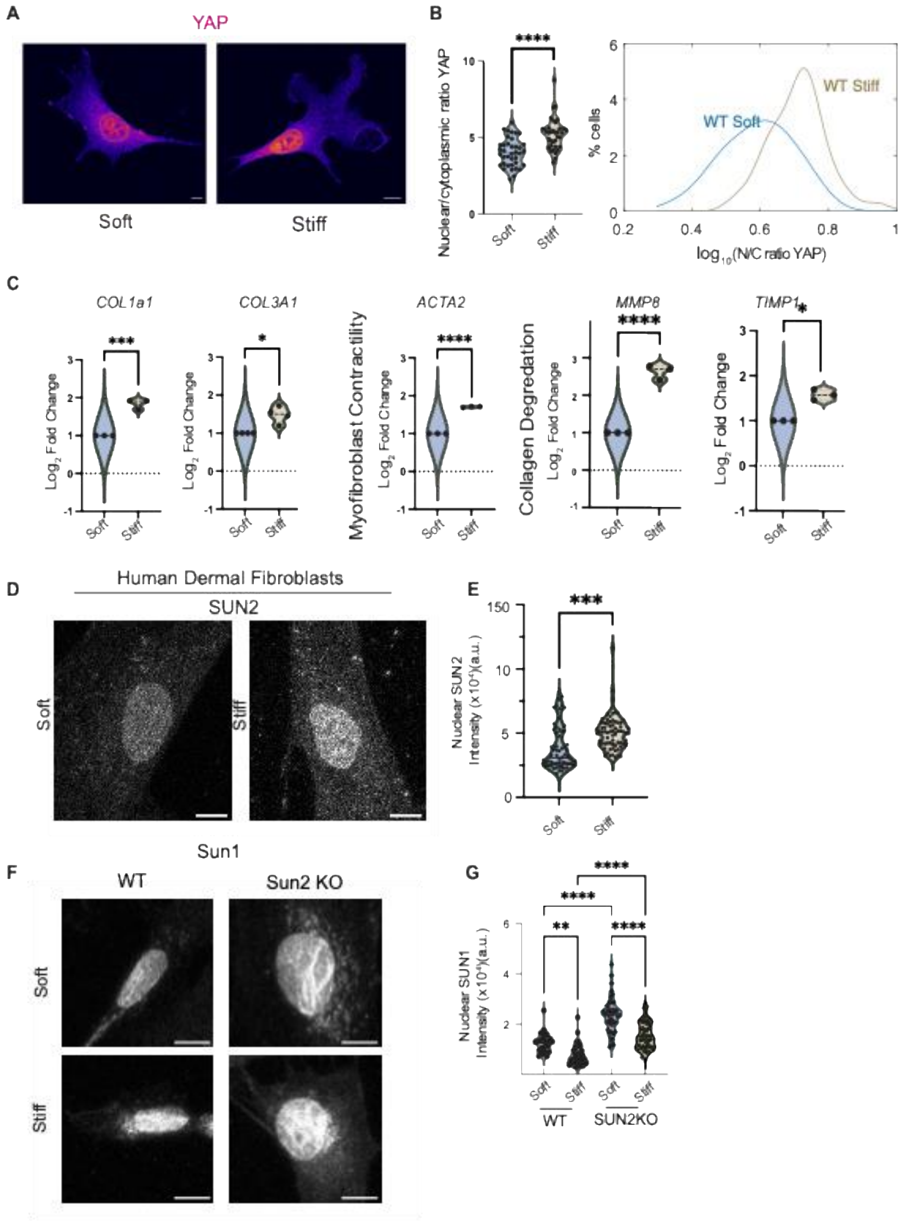


***Figure S1*. Mechanosensing of mouse and human fibroblasts.** *(A) Representative images of YAP localization on soft versus stiff substrates. (B) Quantification of YAP nuclear-to-cytoplasmic (N/C) ratio (left) and distribution of* $\log_{10}$*(N/C ratio) values (right).* $N = 3$, $n_{soft} = 33$, $n_{stiff} = 20$. *(C) qPCR of the indicated genes (COL1A1, COL3A1, ACTA2, MMP8, TIMP1) shown as* $\log_2$ *fold change.* $N = 3$ *independent experiments. (D) Representative immunofluorescence images of SUN2 in human dermal fibroblasts cultured on soft versus stiff substrates. (E) Quantification of nuclear SUN2 intensity in human dermal fibroblasts.* $N = 2$, $n_{soft} = 60$, $n_{stiff} = 69$ *(F) Representative images of SUN1 in WT and SUN2 KO fibroblasts cultured on soft or stiff substrates. (G) Quantification of nuclear SUN1 intensity across genotype and stiffness conditions.* $N = 2$, $n_{soft} = 26|43$, $n_{stiff} = 34|39$. *Scale bars= 10 µm. Violin plots show the median and quartiles; each point represents a single cell.*

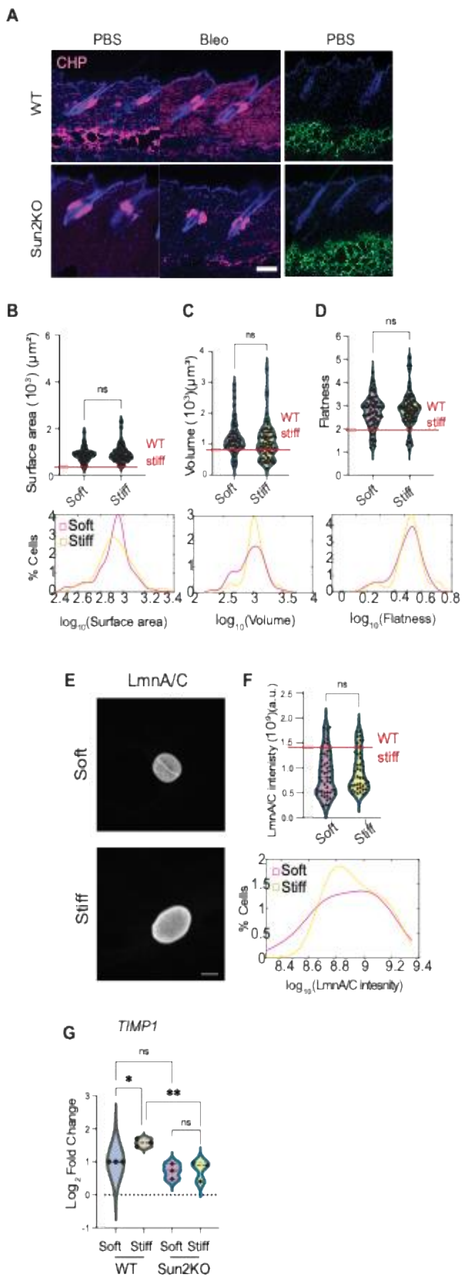


2

***Figure S2. Fibrosis and mechanosensing of dermal fibroblasts.*** *(A) Representative mouse skin sections from PBS- and bleomycin-treated animals stained with collagen hybridizing peptide (CHP; magenta) and DAPI (blue); right, representative adipose layer staining (green) with DAPI (blue). (B–D) Violin plots (top) and corresponding distributions (bottom) of the 3D nuclear morphology of WT mouse fibroblasts on soft versus stiff substrates; ns, not significant. (E) Representative immunofluorescence images of LmnA/C in WT fibroblasts cultured on soft or stiff substrates. (F) Right, violin plot of mean nuclear LMNA/C intensity (top) and probability distribution of* $log_{10}$ *(LmnA/C intensity) (bottom) for murine SUN2 KO cells on soft (pink) and stiff (yellow) substrates, red line denotes mean levels on WT fibroblasts on stiff substrates; ns, not significant.* $N = 3$, $n_{soft} = 49$, $n_{stiff} = 45$. *Significance was determined using a two-tailed Mann–Whitney U test. (G) Timp1 expression (*$log_2$ *fold change) in WT and SUN2KO murine fibroblasts cultured on soft or stiff substrates.* $N = 3$ *independent experiments. Significance determined by one way ANOVA.*

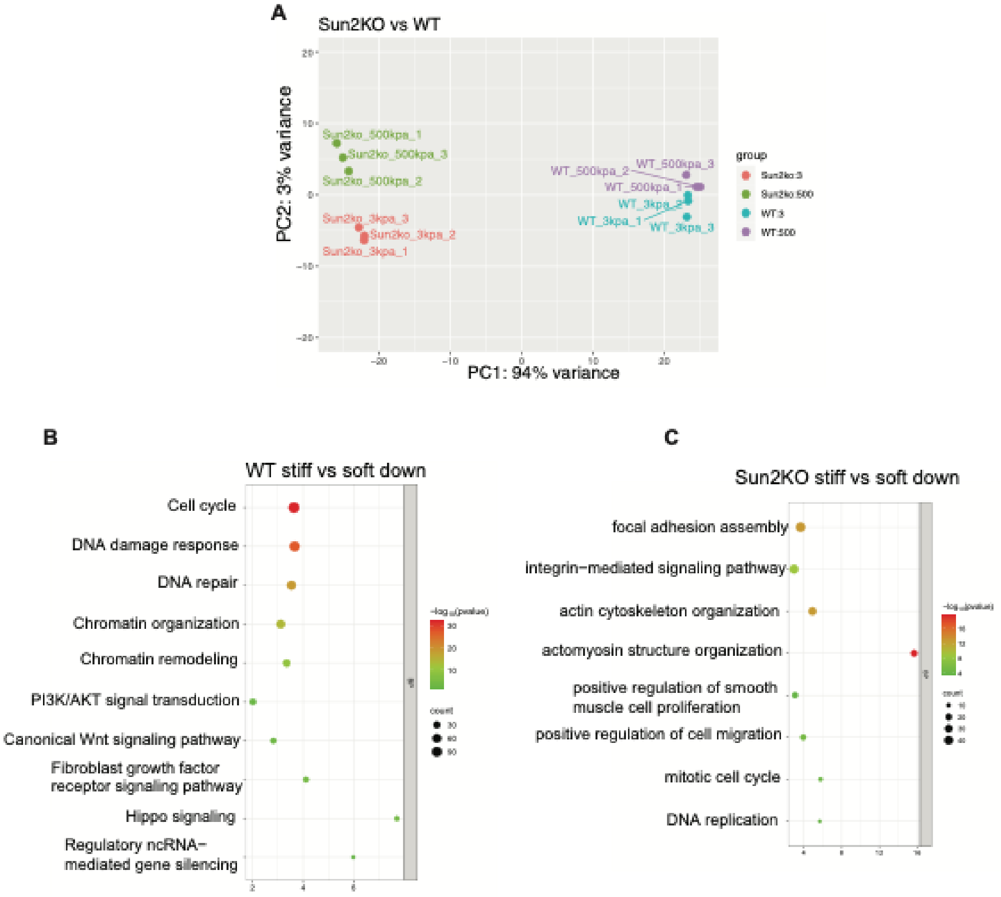


***Figure S3. Analysis of transcriptional changes in Sun2 KO vs WT fibroblasts*** *(A) Principal component analysis (PCA) of bulk RNA-seq samples from WT and SUN2 KO fibroblasts cultured on soft (3 kPa) or stiff (300 kPa). (B–C) Gene Ontology biological process (BP) enrichment of genes downregulated on stiff versus soft substrates in WT (B) or SUN2 KO (C) fibroblasts. Dot position indicates enrichment significance (*$log_{10}$*(p-value)), and dot size denotes the number of genes in each term.*
